# CaaX-like protease of cyanobacterial origin is required for complex plastid biogenesis in malaria parasites

**DOI:** 10.1101/2020.06.02.130229

**Authors:** Thomas R. Meister, Yong Tang, Michael J. Pulkoski-Gross, Ellen Yeh

## Abstract

*Plasmodium* parasites and related apicomplexans contain an essential “complex plastid” organelle of secondary endosymbiotic origin, the apicoplast. Biogenesis of this complex plastid poses a unique challenge requiring evolution of new cellular machinery. We previously conducted a mutagenesis screen for essential apicoplast biogenesis genes to discover organellar pathways with evolutionary and biomedical significance. Here we validate and characterize a gene candidate from our screen, Pf3D7_0913500. Using a conditional knockdown strain, we show that Pf3D7_0913500 depletion causes growth inhibition that is rescued by the sole essential product of the apicoplast, isopentenyl pyrophosphate (IPP), and results in apicoplast loss. Because Pf3D7_0913500 had no previous functional annotation, we name it <u>a</u>picoplast-<u>m</u>inus IPP-<u>r</u>escued 4 (AMR4). AMR4 has an annotated CaaX Protease and Bacteriocin Processing (CPBP) domain, which in eukaryotes typically indicates a role in CaaX post-prenylation processing. Indeed, AMR4 is the only CaaX-like protease in *Plasmodium* parasites which are known to require protein prenylation, and we confirm that the conserved catalytic residue of AMR4 is required for its apicoplast function. However, we unexpectedly find that AMR4 does not act in a CaaX post-prenylation processing pathway in *P. falciparum*. Instead, we find that AMR4 is imported into the apicoplast and is derived from a cyanobacterial CPBP gene which was retained through both primary and secondary endosymbiosis. Our findings suggest that AMR4 is not a true CaaX protease, but instead acts in a conserved, uncharacterized chloroplast pathway that has been retained for complex plastid biogenesis.

**Importance:** *Plasmodium* parasites, which cause malaria, and related apicomplexans are important human and veterinary pathogens. These parasites represent a highly divergent and understudied branch of eukaryotes, and as such often defy the expectations set by model organisms. One striking example of unique apicomplexan biology is the apicoplast, an essential but non-photosynthetic plastid derived from an unusual secondary (eukaryote-eukaryote) endosymbiosis. Endosymbioses are a major driver of cellular innovation, and apicoplast biogenesis pathways represent a hotspot for molecular evolution. We previously conducted an unbiased screen for apicoplast biogenesis genes in *P. falciparum* to uncover these essential and innovative pathways. Here, we validate a novel gene candidate from our screen and show that its role in apicoplast biogenesis does not match its functional annotation predicted by model eukaryotes. Our findings suggest that an uncharacterized chloroplast maintenance pathway has been reused for complex plastid biogenesis in this divergent branch of pathogens.

## Introduction

*Plasmodium spp*., which cause malaria, and related apicomplexans are important human and veterinary pathogens. Beyond their biomedical significance, these parasitic protozoa represent a highly divergent and understudied branch of the eukaryotic tree, distinct from the well-studied model organisms in the opisthokont clade (e.g. yeast, mammals). As such, apicomplexans often defy the expectations of model eukaryotic biology, revealing surprising innovations that both highlight the diversity of eukaryotic life and can be leveraged for therapeutic intervention. One striking illustration of this unique biology is the non-photosynthetic <u>apico</u>mplexan <u>plast</u>id, or apicoplast^1^. The apicoplast is an example of a “complex plastid” derived from secondary endosymbiosis, in which an algae bearing a primary chloroplast was itself engulfed by another eukaryote^2^. Although the apicoplast is no longer photosynthetic, it retains several metabolic pathways, is essential for parasite survival during human infection, and is a proven target of antiparasitic drugs^3–5^.

Despite its importance to pathogenesis, little is known about how the apicoplast is maintained and replicated during the *Plasmodium* life cycle. Like other endosymbiotic organelles, the apicoplast cannot be formed *de novo* and must be faithfully inherited by its growth, division, and segregation into daughter parasites. The few details we know about these apicoplast biogenesis pathways reveal surprising innovations in eukaryotic evolution, particularly in membrane biology^6^. Chloroplasts and endomembranes became uniquely intertwined during secondary endosymbiosis, which enclosed a double-membraned chloroplast in two outer membranes derived from the cell membrane of the algal symbiont and the phagocytic membrane of the engulfing eukaryote. Altogether, the apicoplast is surrounded by four membranes and is, in essence, an endosymbiotic organelle residing within a new membrane compartment. Transit of nuclear-encoded proteins to the apicoplast requires vesicle trafficking from the endoplasmic reticulum (ER) and a unique <u>s</u>ymbiont-derived <u>E</u>R-associated degradation-<u>l</u>ike <u>ma</u>chinery (SELMA) to cross the two outer membranes^7–10^. Additionally, several members of the highly-conserved eukaryotic autophagy pathway are unexpectedly required for apicoplast inheritance^11–14^. While the repurposed SELMA and autophagy components hint at an abundance of molecular innovation in apicoplast biogenesis pathways, most of these pathways remain undiscovered.

To discover new genes required for apicoplast biogenesis and uncover more instances of molecular innovation, we previously conducted a mutagenesis screen to identify mutations that cause apicoplast loss^14^. From this screen, we identified a mutation in a gene encoding a CaaX Protease and Bacteriocin Processing (CPBP) family protein (Pf3D7_0913500, S347G). Known eukaryotic CPBP proteins that have been functionally characterized are CaaX proteases in the post-prenylation processing pathway. During protein prenylation, a hydrophobic prenyl group is covalently attached to the cysteine residue of a C-terminal CaaX motif (where *a* is aliphatic and *X* is any amino acid). Further modification of the prenylated CaaX by the post-prenylation processing pathway includes cleavage of the terminal -aaX by a CaaX protease and carboxyl methylation of the neo C-terminus by an isoprenylcysteine carboxyl methyltransferase (ICMT). Together, these post-translational modifications mediate membrane association and regulate the function of many proteins^15,16^.

Protein prenylation is essential in *P. falciparum*^17,18^ suggesting the presence of a post-prenylation processing pathway. It seemed likely that Pf3D7_0913500 would function as a CaaX protease in this pathway since it is the only *P. falciparum* gene with an annotation consistent with this function. The identification of Pf3D7_0913500 in our apicoplast biogenesis screen prompted several questions: 1) Is Pf3D7_0913500 essential specifically for apicoplast biogenesis? 2) Does Pf3D7_0913500 function as a CaaX protease? 3) Is CaaX post-prenylation processing an apicoplast biogenesis pathway? To answer these questions, we utilized both reverse genetics and molecular evolutionary comparisons to probe the cellular role of Pf3D7_0913500.

## Results

### AMR4 is essential and required specifically for apicoplast biogenesis

We previously performed a mutagenesis screen to identify new apicoplast biogenesis genes^14^. Loss-of-function mutations in these genes disrupt inheritance of the apicoplast during parasite replication resulting in organelle loss in daughter cells. 51 mutants showing apicoplast loss were isolated and analyzed by whole-genome sequencing. To complete our analysis, we sequenced six additional mutant clones from this screen. Two of these clones contained previously identified mutations. We also report missense mutations in two new gene candidates, Pf3D7_1313400 and Pf3D7_1024900 (Table S1). Altogether, out of 57 mutant clones analyzed, we identified 14 gene candidates. In our initial report, we validated three new apicoplast biogenesis genes that contained nonsense mutations. To continue validation of gene candidates identified by this screen, we next focused on genes containing missense mutations. Pf3D7_0913500 was identified by a mutation resulting in an S347G variant but has no known function in apicoplast biogenesis. However, it is predicted to be essential in blood-stage *P. falciparum*^19^ and to contain an apicoplast targeting sequence^20,21^ (Fig. 1A), suggesting that it is an essential apicoplast protein consistent with a role in the organelle’s biogenesis.

**Figure 1.**
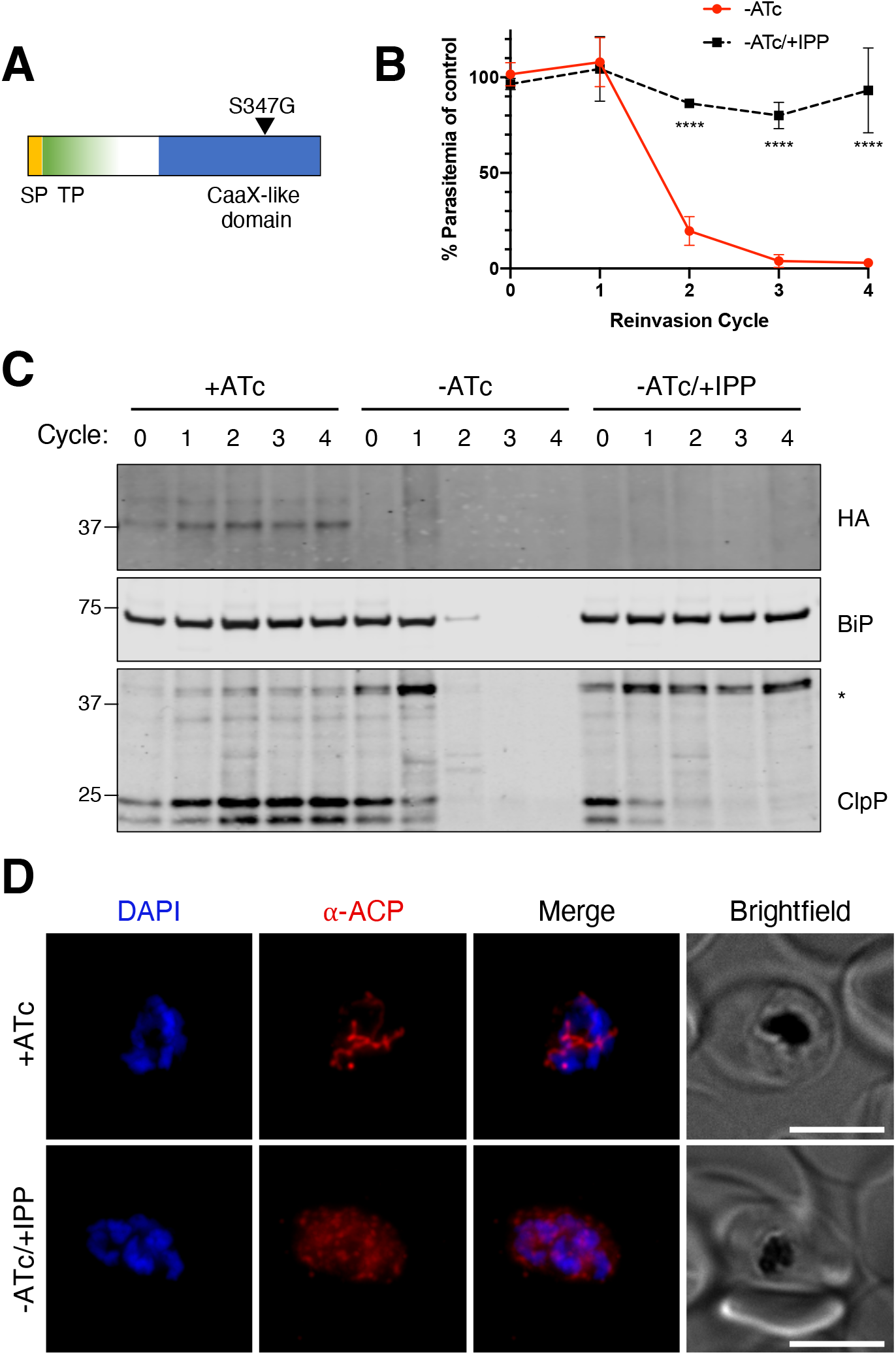
AMR4 is an essential protein required for apicoplast biogenesis. **A.** Domain organization of AMR4 (Pf3D7_0913500) showing predicted signal peptide (SP), apicoplast transit peptide (TP), and the identified mutation. **B.** Growth time course of AMR4 TetR-DOZI knockdown parasites in the absence of ATc with and without IPP. Data are normalized to +ATc control at each time point. Error bars represent standard deviation of the mean of 3 biological replicates. **** *P* < 0.0001 compared to -ATc condition, repeated measures two-way ANOVA with Tukey’s multiple comparisons test. **C.** Western blot from growth time course of AMR4 knockdown parasites showing expression of AMR4-3xHA and processing of ClpP. Full-length ClpP (marked with asterisk) is ~40 kDa, while processed ClpP after removal of its transit peptide is ~25 kDa. BiP serves as a loading control and marker for parasite growth. Blots are representative of 3 biological replicates. **D.** Representative immunofluorescence images showing localization of the apicoplast marker ACP in +ATc and −ATc/+IPP parasites at reinvasion cycle 4. Scale bars 5μm.

To test the essentiality of Pf3D7_0913500, we generated a conditional knockdown strain by modifying the endogenous locus to include a C-terminal triple hemagglutinin (HA) tag and a 3’ UTR RNA aptamer sequence that binds a tetracycline repressor (TetR) and development of zygote inhibited (DOZI) fusion protein^22,23^. In the presence of anhydrotetracycline (ATc), the 3’ UTR aptamer is unbound and Pf3D7_0913500 is expressed. Removal of ATc from the media early in the parasite replication cycle resulted in efficient knockdown of Pf3D7_0913500 within the same cycle (Fig. 1C). We monitored parasite growth over several reinvasion cycles and found that depletion of Pf3D7_0913500 caused significant growth inhibition (Fig. 1B), confirming its essentiality. The only essential product of the apicoplast in blood-stage *Plasmodium* is isopentenyl pyrophosphate (IPP), such that supplementation with exogenous IPP rescues parasites with apicoplast defects including complete loss of the organelle^24^. Consistent with Pf3D7_0913500 having a specific apicoplast function, growth inhibition caused by Pf3D7_0913500 knockdown was fully reversed with the addition of IPP (Fig. 1B).

Essential apicoplast functions fall into two broad categories: those involved in organelle biogenesis and those involved solely in IPP production. As indicated, disruption of genes required for organelle biogenesis leads to apicoplast loss; in contrast, disruption of genes only involved in IPP production does not^25^. To distinguish whether Pf3D7_0913500 knockdown caused apicoplast loss, we monitored apicoplast structure by immunofluorescence. During knockdown in IPP-rescued parasites, the apicoplast marker acyl-carrier protein (ACP) redistributed from a single organellar structure to diffuse puncta (Fig. 1D) as previously described for apicoplast loss. We additionally monitored apicoplast protein import as a readout for organelle loss. Most apicoplast proteins have an N-terminal transit peptide that is cleaved upon apicoplast import, and this processing event is eliminated in apicoplast-minus parasites^24^. During Pf3D7_0913500 knockdown in IPP-rescued parasites, we observed the accumulation of the apicoplast protein caseinolytic protease P (ClpP) at its unprocessed molecular weight (Fig. 1C), indicating a disruption in apicoplast biogenesis processes^26^. Together, these results confirm that Pf3D7_0913500 is specifically essential for an apicoplast biogenesis function. Because the gene had no previous functional annotation, we name it <u>a</u>picoplast-<u>m</u>inus IPP-<u>r</u>escued 4 (AMR4), consistent with the loss-of-function phenotype observed in previously validated candidates from our screen.

### A conserved catalytic residue in the protease domain of AMR4 is required for its apicoplast function

The AMR4 mutation identified in our screen (S347G) is near the active site of the gene’s only annotated domain, a CaaX Protease and Bacteriocin Processing (CPBP) domain. Mutagenesis and structural studies of CPBP domains indicate a catalytic mechanism in which a conserved glutamate activates a nucleophilic water molecule for proteolysis^27–30^. Mutation of this catalytic glutamate was shown to completely abolish protease function without disrupting the stability or conformation of the protein^29^. The catalytic glutamate is conserved in AMR4 at position 352. To test whether E352 is required for AMR4 function, we complemented the AMR4 knockdown strain with an episome expressing either wild-type (AMR4^wt^) or protease-dead (AMR4^E352A^) constructs^31^. Upon downregulation of endogenous AMR4 expression, we found that episomal expression of AMR4^wt^ partially but significantly complements the knockdown (Fig. 2A-B, S1). In contrast, episomal expression of AMR4^E352A^ shows no rescue, suggesting that the E352 residue is required for AMR4 function.

**Figure 2.**
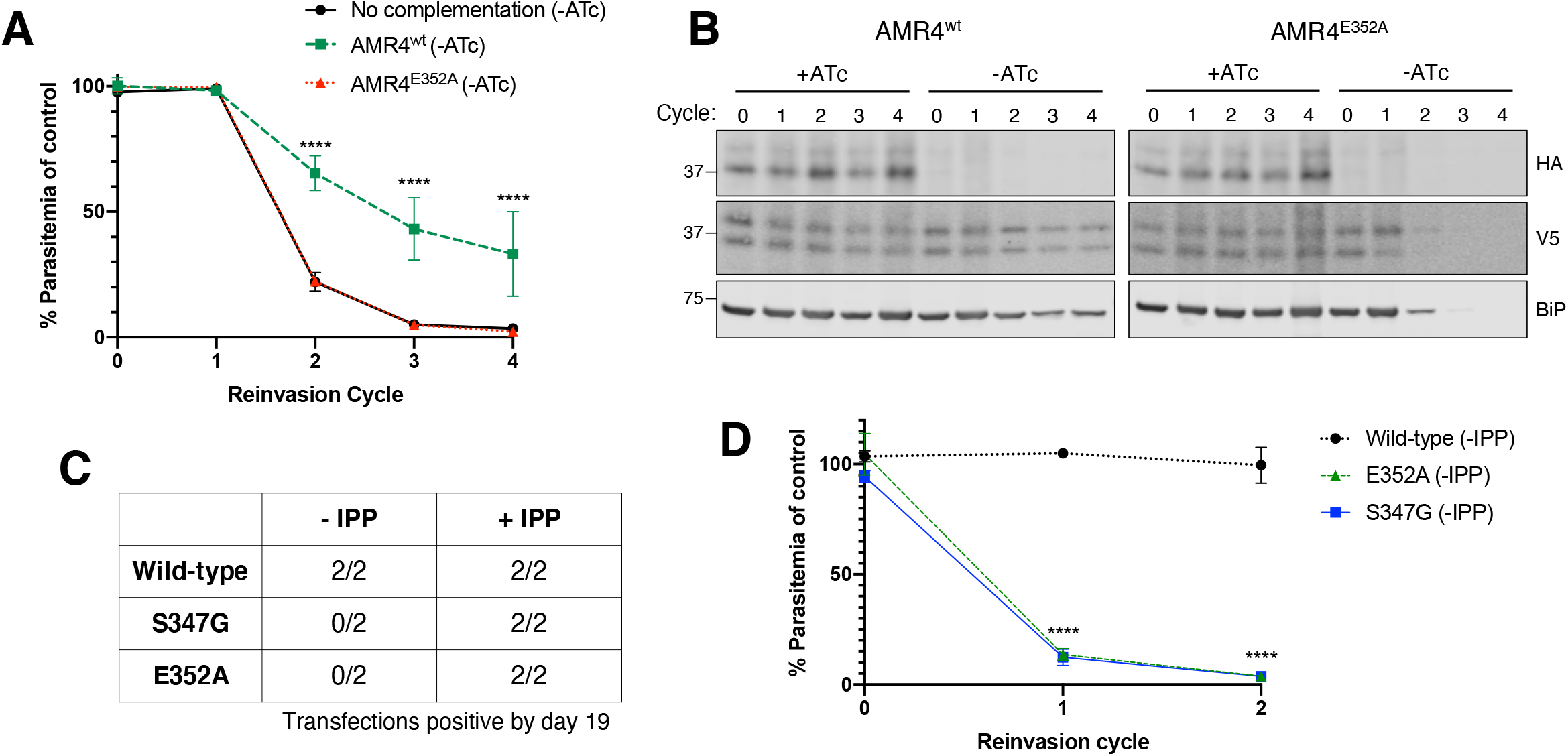
A conserved catalytic residue in the protease domain of AMR4 is required for its apicoplast function. **A.** Growth time course of AMR4 TetR-DOZI knockdown parasites in the absence of ATc, complemented with either AMR4^wt^, AMR4^E352A^, or no complementation construct. Data are normalized to +ATc controls at each time point. Error bars represent standard deviation of the mean of 3 biological replicates. **** P < 0.0001 compared to no complementation control, repeated measures two-way ANOVA with Tukey’s multiple comparisons test. **B.** Western blot from growth time course of AMR4-3xHA knockdown parasites complemented with V5-tagged AMR4^wt^ or AMR4^E352A^. BiP serves as a loading control and marker for parasite growth. **C.** Recovery of transfectants from endogenous AMR4 knock-in of wild-type, S347G, or E352A alleles. Transfections were performed with and without IPP in 2 independent experiments, and growth was monitored by luciferase signal. **D.** Growth time course after removal of IPP from AMR4 knock-in strains recovered under IPP supplementation. Data are normalized to +IPP control at each time point. Error bars represent standard deviation of the mean of 2 biological replicates. **** P < 0.0001 compared to wild-type control, repeated measures two-way ANOVA with Tukey’s multiple comparisons test.

It is likely that the partial functional complementation by AMR4^wt^ is due to altered abundance or stage-dependent expression from the non-native promoter used for episomal expression. As an alternative to episomal expression, we directly introduced the E352A mutation or wild-type control into the endogenous AMR4 locus. We also introduced the S347G mutation identified in our screen to determine whether this mutation disrupted AMR4 function. These “knock-in” transfections were performed with and without IPP supplementation in two independent experiments. Wild-type transfectants were recovered from every transfection, while E352A and S347G transfectants were only recovered with IPP supplementation (Fig. 2C). For transfectants recovered with IPP supplementation, we then removed IPP from the media and tracked growth for two cycles. The E352A and S347G mutants had significantly reduced growth, indicating a dependence on exogenous IPP due to disruption of the apicoplast (Fig. 2D). In contrast, wild-type parasites grew similarly in the presence or absence of IPP supplementation, confirming that the IPP dependence of the E352A and S347G mutants was a result of the mutation and not prolonged growth with IPP supplementation. Together, these results show that the catalytic E352 residue in the CPBP domain is required for AMR4’s apicoplast function, indicating that AMR4 likely retains a protease activity. It additionally validates that the S347G mutation, which identified AMR4 in our initial screen, disrupts the protein’s function and caused the apicoplast loss phenotype in the original mutant.

The only known catalytic activity of eukaryotic CPBP domains is cleavage of a C-terminal CaaX motif (where *a* is aliphatic and *X* is any amino acid) following cysteine prenylation as part of a post-prenylation processing pathway^15,16^. Because CPBP catalysis is required for AMR4 function, we tested whether AMR4 has CaaX protease activity in yeast by functional complementation of a CaaX protease-knockout strain^32^. Although AMR4 did not restore CaaX protease activity in the knockout yeast strain (Fig. S2), we were unable to detect tagged AMR4 to confirm protein expression and proper localization. Therefore, it remains unclear whether AMR4 retains CaaX protease activity.

### AMR4 is an imported apicoplast protein that does not share a post-prenylation processing pathway with *Pf*ICMT

We also sought evidence that AMR4 might participate in a post-prenylation processing pathway in *Plasmodium* parasites. Protein prenylation has been shown to be essential in *P. falciparum*^17,18^. It is presumed that *Plasmodium*, like other organisms, contains a two-step post-prenylation processing pathway, in which a CaaX protease removes the terminal -aaX from prenylated proteins and the neo C-terminus is then methylated by an isoprenylcysteine carboxyl methyltransferase (ICMT)^16^. Two types of eukaryotic CaaX proteases are known: Rce1 glutamic proteases in the CPBP family and Ste24 zinc metalloproteases which are unrelated to the CPBP family. Unlike other apicomplexans and related chromerids, the only annotated CaaX-like protease in *Plasmodium spp.* is AMR4 (Fig. 3A). *Plasmodium spp.* also retain an ICMT homolog, suggesting that AMR4 and *Pf*ICMT may function together in a post-prenylation processing pathway that is essential for apicoplast biogenesis.

**Figure 3:**
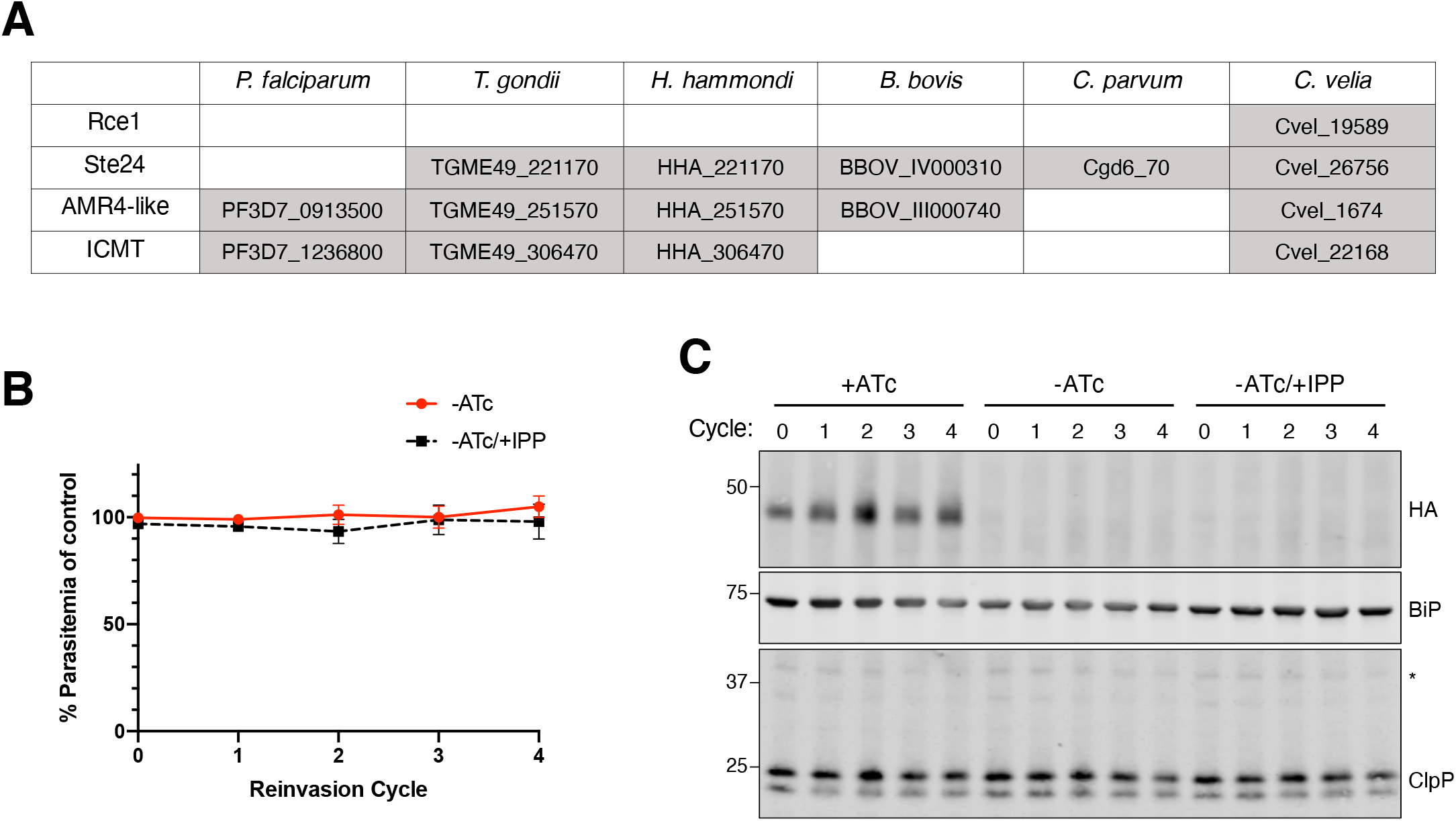
*Pf*ICMT is not an essential gene and does not have an apicoplast function. **A.** Table showing the presence or absence of Rce1, Ste24, AMR4, and ICMT homologs in apicomplexans and chromerids. **B.** Growth time course of *Pf*ICMT TetR-DOZI knockdown parasites in the absence of ATc with and without IPP. Data are normalized to +ATc control at each time point. Error bars represent standard deviation of the mean of 3 biological replicates. No significant differences between means were detected at any time point, repeated measures two-way ANOVA with Tukey’s multiple comparisons test. **C.** Western blot from growth time course of *Pf*ICMT knockdown parasites showing expression of *Pf*ICMT-3xHA and processing of ClpP. Full-length ClpP (marked with asterisk) is ~40 kDa, while processed ClpP after removal of its transit peptide is ~25 kDa. BiP serves as a loading control and marker for parasite growth. Blots are representative of 3 biological replicates.

To determine if *Pf*ICMT is essential and required for apicoplast biogenesis, we generated a conditional knockdown strain using the TetR-DOZI system^22,23^. As expected, removal of ATc caused a significant decrease in protein levels within a single replication cycle (Fig. 3C). However, *Pf*ICMT knockdown did not affect parasite growth over four replication cycles (Fig. 3B), indicating that it is non-essential in blood-stage culture. We also could not detect any defect in processing of the apicoplast protein ClpP (Fig. 3C), confirming that apicoplast biogenesis was not disrupted by *Pf*ICMT knockdown. The dispensability of *Pf*ICMT was surprising since it is proposed to act in an essential post-translational modification pathway. It is possible that our knockdown was insufficient to completely disrupt *Pf*ICMT function, however our result is also supported by an insertional transposon mutagenesis screen which assigns *Pf*ICMT as dispensable^19^.

The post-prenylation processing enzymes typically colocalize to the ER membrane^33–35^. We reasoned that if AMR4 and *Pf*ICMT perform post-prenylation processing together, they should localize to the same membrane. To determine the cellular localization of AMR4 and *Pf*ICMT, we performed immunofluorescence and transit peptide cleavage assays with the endogenously tagged TetR-DOZI strains. Consistent with its predicted apicoplast targeting sequence, co-IFA shows that AMR4 co-localizes with the apicoplast marker ACP and is present on the organelle’s distinctive branching structures during schizogyny (Fig. 4A). When parasites were treated with an inhibitor (actinonin) that causes apicoplast loss, ACP and AMR4 both redistribute to diffuse puncta, as previously observed for known apicoplast proteins^24^. In contrast, *Pf*ICMT does not colocalize with ACP, and its localization is unaffected by actinonin treatment (Fig. 4B), indicating that it does not localize to the apicoplast.

**Figure 4.**
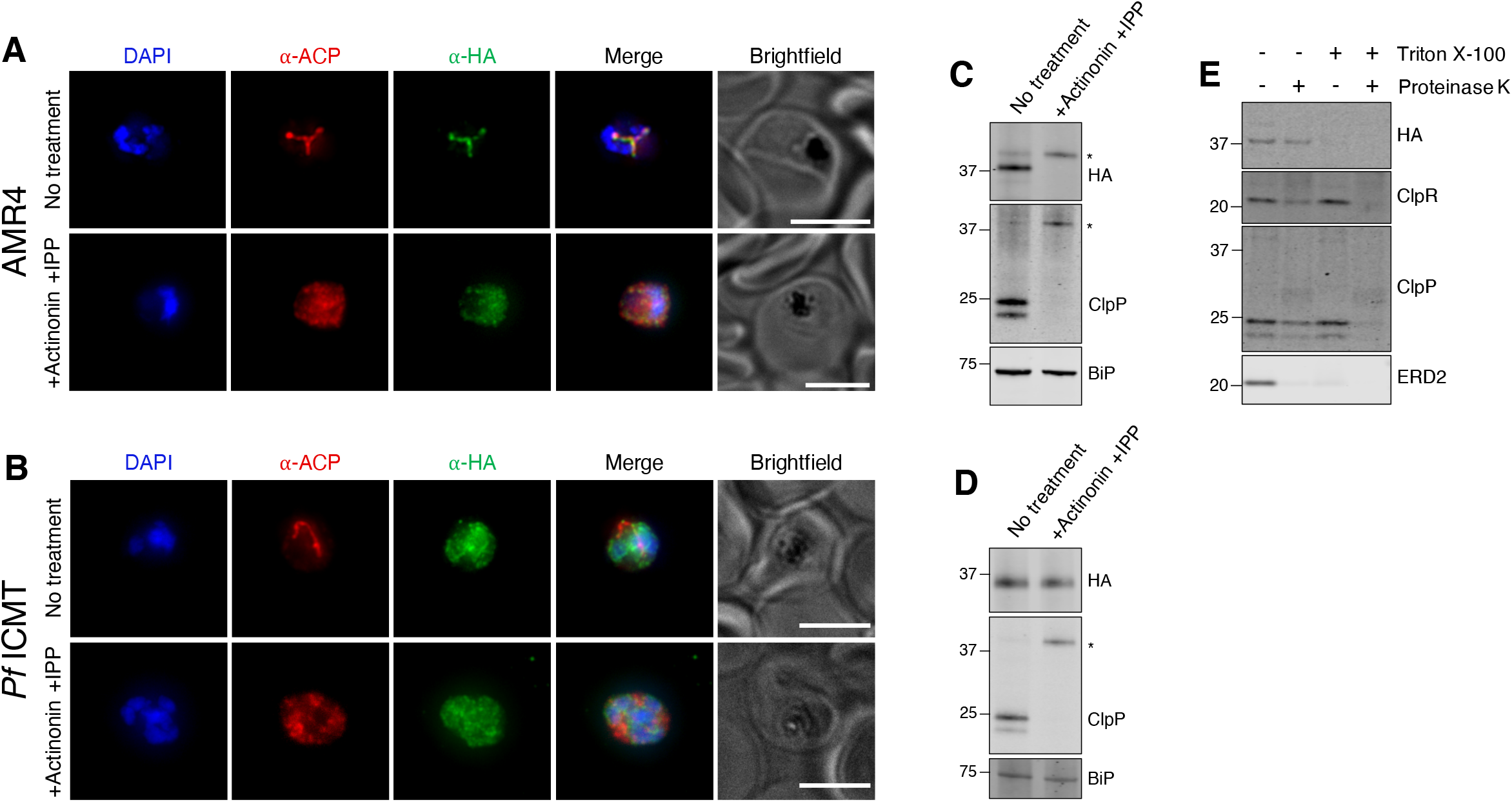
AMR4 is an imported apicoplast protein and does not share a post-prenylation processing pathway with *Pf*ICMT. **A and B.** Representative immunofluorescence images showing localization of AMR4-3xHA (A) and *Pf*ICMT-3xHA (B) compared to the apicoplast marker ACP after 3 days of either no treatment or +10 μM actinonin/200 μM IPP. Scale bars 5μm. **C and D.** Western blots showing processing of AMR4-3xHA (C), *Pf*ICMT-3xHA (D), and ClpP (C and D) after 3 days of either no treatment or +10 μM actinonin/200 μM IPP. BiP serves as a loading control. Blots are representative of 3 biological replicates. **E.** Western blot from protease protection assay. Organellar fraction from AMR4-3xHA parasites was treated with or without Proteinase K in the presence or absence of Triton X-100. ClpR and ClpP serve as controls for internal apicoplast proteins, while ERD2 serves as a control for cytosol-exposed membrane proteins.

We next assessed AMR4 and *Pf*ICMT protein cleavage as a marker for apicoplast import. Most apicoplast proteins have an N-terminal transit peptide that is cleaved upon apicoplast import, and this processing is eliminated in apicoplast-minus parasites^24^. When the apicoplast is intact, AMR4 is detectable as a predominant mature band and a less abundant unprocessed band which likely indicates protein that is en route to the apicoplast. Upon apicoplast loss, AMR4 shifts exclusively to its unprocessed form (Fig. 4C), indicating that it is subject to apicoplast-dependent N-terminal cleavage. In contrast, *Pf*ICMT is detected as a single band which is unaffected by actinonin treatment (Fig. 4D), consistent with its non-apicoplast localization.

CaaX prenylation occurs in the cytoplasm, followed by post-prenylation processing on a cytosolic membrane face. We reasoned that for AMR4 to perform a CaaX protease function, it should localize to the outer apicoplast membrane. To test this, we performed a protease protection assay in which we hypotonically lysed cell membranes of the endogenously tagged AMR4 strain while leaving organellar membranes intact. We then incubated this sample with Proteinase K and assessed proteolysis by western blot. As expected, the mature forms of internal apicoplast proteins ClpP and ClpR are largely resistant to proteolysis (42 and 47%, respectively). Similarly, the mature form of AMR4 is resistant to proteolysis (74%), indicating that it is imported from the outer membrane of the apicoplast (Fig. 4E). In contrast, the cytosol-exposed ER/Golgi membrane protein ERD2^36^ is completely proteolyzed. When Proteinase K is added in the presence of 1% Triton X-100 to disrupt organellar membranes, both cytosol-exposed and membrane-protected proteins are proteolyzed. We note that the integral membrane proteins ERD2 and AMR4 are also degraded after Triton lysis in absence of Proteinase K, likely indicating that the membrane-extracted proteins are exposed to endogenous proteases during incubation. Based on these results, we conclude that AMR4 is localized to the apicoplast and is imported into one of the inner membranes, unlike typical eukaryotic CaaX proteases. Additionally, *Pf*ICMT does not phenocopy or colocalize with AMR4, suggesting that they do not share a post-prenylation processing pathway.

### AMR4 is derived from a prokaryotic CPBP gene acquired through endosymbiosis

The CPBP family consists of two branches: eukaryotic Rce1 CaaX proteases and prokaryotic abortive infection (abi) proteases. Because AMR4 is apicoplast-localized and does not function in a CaaX post-prenylation processing pathway, we hypothesize that it may have originated from the prokaryotic chloroplast-like symbiont and not from a eukaryotic Rce1 from either the red algal symbiont or secondary host. To determine its origin, we generated a phylogeny of CPBP proteins from organisms along each step of these successive endosymbioses, including 1) cyanobacteria 2) eukaryotes without plastids 3) eukaryotes with primary chloroplasts 4) eukaryotes with secondary red plastids. AMR4 is highly conserved in apicomplexans and related chromerids, with the exception of *Cryptosporidium* which has lost its apicoplast. The phylogeny shows with strong support that AMR4 and its secondary plastid homologs form a monophyletic clade with both cyanobacterial proteins and primary chloroplast proteins with predicted transit peptides, including *At*Sco4 which is known to be chloroplast-localized^37^ (Fig. 5, Table S2). Outside of this clade are eukaryotic Rce1’s which do not have predicted chloroplast transit peptides, including confirmed ER CaaX proteases from yeast and *Arabidopsis*^33,34^. To prevent bias from different targeting signals in our analysis, we also generated a phylogeny using only the annotated CPBP domain (IPR003675) from each protein, which confirmed AMR4’s inclusion in the cyanobacterial/plastid clade (Fig. S3).

**Figure 5.**
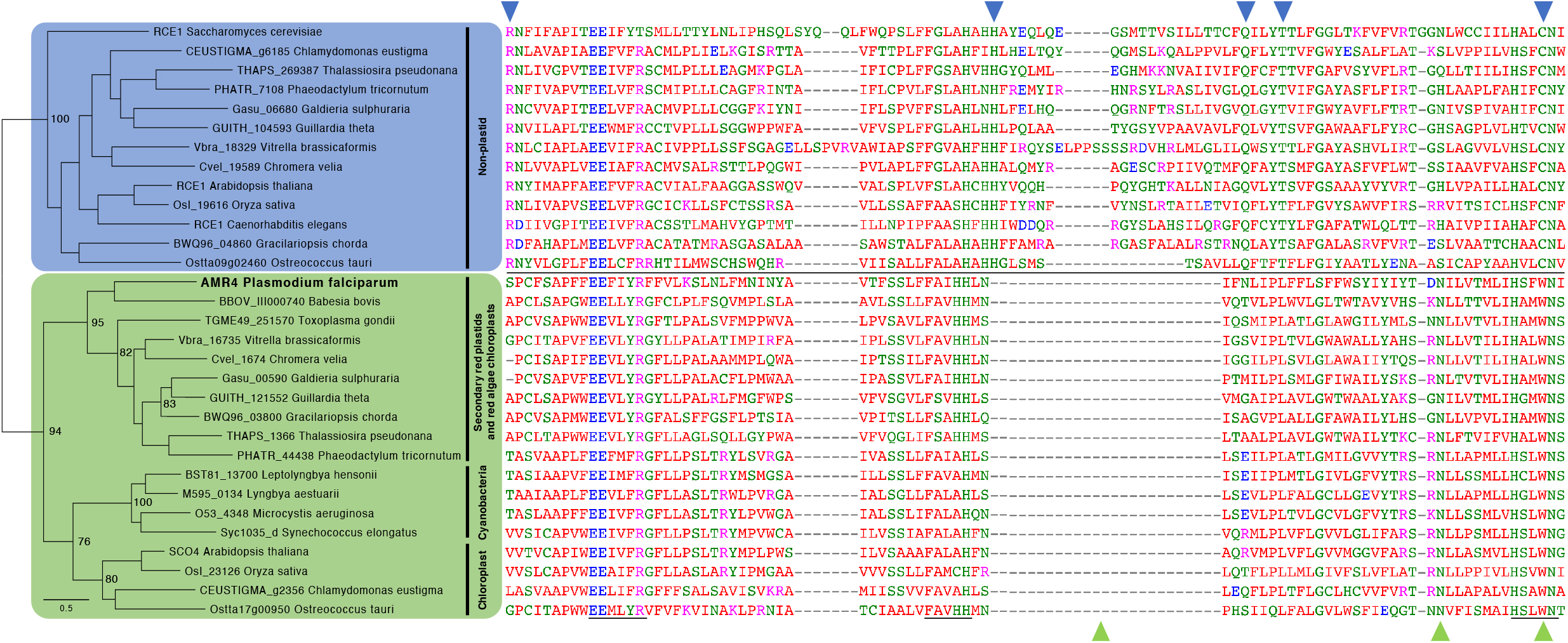
AMR4 is derived from a prokaryotic CPBP gene acquired through endosymbiosis. Phylogenetic analysis of selected CPBP proteins from cyanobacteria, primary and secondary plastids, and non-plastid Rce1’s. Maximum likelihood phylogeny defines a monophyletic clade of cyanobacterial and chloroplast-targeted proteins along with AMR4 and its secondary plastid homologs. Outside of this clade are non-plastid targeted Rce1 proteins from eukaryotes with and without plastids. Branch support values for well-supported major nodes are shown, out of 100 bootstrap intervals. Sequence alignment of the CPBP domain from each protein is shown. Arrowheads indicate residues which are conserved among non-plastid Rce1’s (blue) or among cyanobacterial and plastid CPBP’s (green). Conserved CPBP motifs are underlined. Predicted targeting sequences for all proteins are shown in Table S2.

Sequence alignment of these CPBP domains highlights several residues and motifs which are highly conserved within either eukaryotic Rce1’s or cyanobacterial-derived proteins (Fig. 5). In each of these cases, AMR4 shares the conserved sequence with cyanobacterial proteins, but does not share any of the Rce1-specific motifs. Taken together, these results confirm that AMR4 originated from a cyanobacterial CPBP gene which was retained through both primary and secondary endosymbiosis. This further suggests that AMR4 may share a molecular function with its homologs in chloroplasts and other complex plastids.

## Discussion

As a result of its divergent evolution, thousands of genes in the *P. falciparum* genome remain unannotated^38^. We previously designed an unbiased screen that allows for discovery of new genes with essential roles in apicoplast biogenesis^14^. Here, we validate a previously uncharacterized gene candidate from our screen, AMR4, and show that it has an essential function in apicoplast biogenesis. AMR4 has an annotated CaaX Protease and Bacteriocin Processing (CPBP) domain, which usually indicates a CaaX post-prenylation processing function in eukaryotes. However, we provide three lines of evidence to demonstrate that AMR4 does not perform post-prenylation processing. First, we show that the downstream post-prenylation processing enzyme *Pf*ICMT does not phenocopy or colocalize with AMR4, suggesting that they do not share a pathway. Second, we show that AMR4 is imported into one of the inner apicoplast membranes, inconsistent with proteolysis of prenylated CaaX proteins on cytosolic membrane faces. Finally, we find that AMR4 did not evolve from a eukaryotic CaaX protease, but instead is derived from a cyanobacterial CPBP gene which was retained through both primary and secondary endosymbiosis.

Although AMR4 is not a CaaX protease, we show that the catalytic residue of its CPBP domain is required for its role in apicoplast biogenesis. This suggests that AMR4 retains protease activity, however its substrate(s) and pathway partners are unknown. Our finding that AMR4 was derived from a cyanobacterial CPBP gene, as opposed to a eukaryotic CaaX protease from either the red algal endosymbiont or secondary host, suggests that it might exist as part of a conserved chloroplast pathway. Indeed, a chloroplast-targeted CPBP protein in *A. thaliana* (*At*Sco4) was previously shown to play a role in chloroplast maintenance and light acclimation^37^. It is possible that AMR4 and its secondary plastid homologs act in a similar pathway as *At*Sco4. However, many primary chloroplasts including *A. thaliana* have an expanded repertoire of uncharacterized CPBP proteins^39^, therefore limiting our ability to infer AMR4 function. Altogether, we consider it likely that AMR4 resides in one of the inner two apicoplast membranes and performs a conserved but unknown role in plastid maintenance. Future work will focus on identifying substrates of AMR4 proteolysis and uncovering the apicoplast biogenesis pathway in which it functions.

Our findings also raise questions about the state of post-prenylation processing in *Plasmodium*. Do *Plasmodium spp.* possess a functional CaaX protease? AMR4 is the only identifiable CaaX-like protease, however our results show that it does not perform a CaaX function. Therefore, it appears that *Plasmodium spp.* might lack a CaaX protease, in contrast to other apicomplexan parasites which retain a Ste24 homolog that could perform this function (Fig. 3A). Alternatively, *Plasmodium spp.* may possess a cryptic CaaX protease which has not yet been identified, either a highly divergent Ste24 homolog or an unrelated protease which has evolved a post-prenylation processing function.

If *Plasmodium* lacks a CaaX protease, is the post-prenylation processing pathway being lost? Genome reduction is a major driving force during parasite evolution, and therefore non-essential pathways are likely to be lost^40^. While prenylation is known to be essential in *P. falciparum*^17,18^, we show that the post-prenylation processing enzyme *Pf*ICMT is non-essential in blood-stage culture. This is consistent with results from previous whole-genome essentiality screens which assign “dispensable” annotations to *Pf*ICMT as well as the *T. gondii* Ste24 and ICMT homologs^19,41^. Additionally, several apicomplexan genera including *Babesia* and *Cryptosporidium* lack ICMT homologs (Fig. 3A), suggesting partial loss of the post-prenylation processing pathway in related parasites. However, we cannot rule out the possibility that apicomplexan Ste24 or ICMT homologs are required in other parasite life stages and growth conditions or that cryptic ICMT activity may be required.

Do prenylated CaaX proteins undergo further modification in *Plasmodium*? *P. falciparum* has a reduced prenylome, containing only 8 prenylated CaaX proteins^42,43^. If none of these proteins require post-prenylation processing to function, then it is likely that the pathway would be lost. Indeed, previous work in model systems shows that not all prenylated CaaX proteins undergo further modification from post-prenylation processing enzymes^44,45^. Similar proteomic approaches will be required in *Plasmodium* to determine if any prenylated proteins undergo CaaX cleavage and/or carboxyl methylation, indicating whether the parasites retain a cryptic CaaX protease or whether the post-prenylation processing pathway has been lost.

More broadly, our results highlight the need to study eukaryotes that diverge from model organisms. We provide another example of apicomplexan biology defying the expectations set by model eukaryotes. Our screen provides an unbiased method to identify divergent genes that act in essential apicoplast biogenesis pathways. Apicoplast biogenesis is a hotspot for molecular innovation, as demonstrated by examples of genes being reused (e.g. AMR4), repurposed (e.g. Atg8, SELMA, AMR1), or newly invented (e.g. AMR3) to maintain the complex plastid. In each of these cases, gene annotations based on model eukaryotes fail to accurately represent biological function. These novel pathways must be uncovered through biochemical, cellular, and comparative evolutionary approaches to better understand the true diversity of eukaryotic life.

## Materials and Methods

### Ethics statement

Human erythrocytes were purchased from the Stanford Blood Center (Stanford, CA) to support *in vitro* P. falciparum cultures. Because erythrocytes were collected from anonymized donors with no access to identifying information, IRB approval was not required. All consent to participate in research was collected by the Stanford Blood Center.

### Parasite culture and transfections

*Plasmodium falciparum* parasites were grown in human erythrocytes (Stanford Blood Center) at 2% hematocrit in RPMI 1640 media (Gibco) supplemented with 0.25% Albumax II (Gibco), 2 g/L sodium bicarbonate (Fisher), 0.1 mM hypoxanthine (Sigma), 25 mM HEPES, pH 7.4 (Sigma), and 50 μg/L gentamycin (Gold Biotechnology) at 37°C, 5% O_2_, and 5% CO_2_.

Transfections were performed into NF54^Cas9+T7 Polymerase^ parasites^41^ (a kind gift from Jacquin Niles) using variations of the spontaneous uptake method^23,46^. In the first variation, 100 μg of each plasmid was ethanol precipitated and resuspended in 0.2 cm electroporation cuvettes in 30 μL sterile TE buffer, 170 μL cytomix, and 200 μL packed erythrocytes. Erythrocytes were electroporated at 310 V, 950 μF, infinite resistance in a Gene Pulser Xcell electroporation system (Bio-Rad) before allowing parasites to invade. Drug selection was initiated 3 days after transfection. In the second variation, 50 μg of each plasmid was ethanol precipitated and resuspended in 0.2 cm electroporation cuvettes in 100 μL TE buffer, 100 μL RPMI 1640 containing 10 mM HEPES-NaOH, pH 7.4, and 200 μL packed erythrocytes. Erythrocytes were pulsed with 8 square wave pulses of 365 V x 1 ms separated by 0.1 s. Erythrocytes were allowed to reseal for 1 hour in a 37°C water bath before allowing parasites to invade. Drug selection was initiated 4 days after transfection.

All transfectants were selected with 2.5 μg/mL Blasticidin S (Research Products International) and maintained in the presence of 500 nM anhydrotetracycline (ATc, Sigma). Additionally, strains carrying the pfYC110 episome were selected and maintained with 125 μg/mL G418 Sulfate (Corning). Integration of plasmids into endogenous loci was confirmed by PCR. Additionally, integration of the correct allele from each knock-in mutation was confirmed by Sanger sequencing. Growth of knock-in transfections was monitored using the *Renilla* Luciferase Assay System (Promega) and GloMax 20/20 Luminometer (Promega).

### Vector construction

Oligonucleotides and gBlocks were ordered from IDT. Molecular cloning was performed using Gibson Assembly (NEB) or In-Fusion Cloning (Clonetech). All primer and gBlock sequences are provided in Table S3.

For CRISPR-Cas9-based editing of endogenous Pf3D7_0913500 (AMR4) and Pf3D7_1236800 (*Pf*ICMT) loci, sgRNA’s were designed using the eukaryotic CRISPR guide RNA/DNA tool (http://grna.ctegd.uga.edu/). To generate a linear plasmid for CRISPR-Cas9-based editing, left and right homology regions were amplified for each gene, and a gBlock containing the recoded sequence C-terminal of the CRISPR cut site and a triple HA tag was synthesized with appropriate overhangs for Gibson Assembly. This fragment and the left homology region were simultaneously cloned into the FseI/ApaI sites of the linear plasmid pSN054-V5. Next, the appropriate right homology region and a gBlock containing the sgRNA expression cassette were simultaneously cloned into the AscI/I-SceI sites of the resultant vectors to generate each TetR-DOZI plasmid. Plasmids for knock-in mutations of the endogenous Pf3D7_0913500 locus were generated as described above, using C-terminal recoded gBlocks with the intended mutation.

To generate the episomal complementation constructs for Pf3D7_0913500, gBlocks containing the recodonized N-terminus and C-terminus (either wild-type or E352A mutant) of Pf3D7_0913500 were stitched together by overlap extension PCR and cloned into the AvrII/SacII sites of the pfYC110 plasmid^31^ using In-Fusion Cloning.

To generate the mating hormone A-factor 1 (MFA1) expression plasmid, the MFA1 5’UTR, coding sequence, and 3’UTR were amplified from *S. cerevisiae* genomic DNA and cloned into the BamHI/HindIII sites of the pRS316 plasmid^32^ using In-Fusion Cloning. To generate the *Sc*Rce1 complementation construct, the 5’ UTR and *Sc*Rce1 coding sequence were individually amplified. These fragments and a gBlock containing a C-terminal Flag tag and the *Sc*Rce1 3’UTR were cloned into the SacII/HindIII sites of the pRS315 plasmid^32^. To generate Pf3D7_0913500 complementation constructs with the *Sc*Rce1 signal peptide, amino acids 112-432 or 197-432 were amplified from a gBlock containing recodonized Pf3D7_0913500. These fragments and gBlocks containing the *Sc*Rce1 signal peptide with appropriate overhangs were then cloned into the BamHI/PstI sites of the resultant *Sc*Rce1 complementation construct using In-Fusion Cloning.

### Western blotting

Parasites were separated from erythrocytes by lysis in 0.1% saponin for 5 minutes on ice. Parasite pellets were washed twice with ice-cold phosphate-buffered saline (PBS), resuspended in PBS containing 1x NuPAGE LDS sample buffer with 50 mM DTT, and boiled for 10 minutes before separation on NuPAGE gels (Invitrogen). Gels were transferred onto nitrocellulose membranes using the Trans-Blot Turbo system (Bio-Rad). Membranes were blocked in 0.1% Hammarsten casein (Affymetrix) in 0.2X PBS with 0.01% sodium azide. Antibody incubations were performed in a 1:1 mixture of blocking buffer and Tris-Buffered Saline with Tween 20 (TBST; 10 mM Tris, pH 8.0, 150 mM NaCl, 0.25 mM EDTA, 0.05% Tween 20). Blots were incubated with primary antibody for either 1 hour at room temperature or at 4°C overnight using the following dilutions: 1:1,000 mouse-α-HA 2.2.14 (Thermo Fisher 26183), 1:4,000 rabbit-α- *Pf*ClpP and 1:4,000 rabbit-α-*Pf*ClpR^47^ (kind gifts from Walid Houry), 1:20,000 mouse-α-*Py*BiP (a kind gift from Sebastian Mikolajczak and Stefan Kappe), 1:1,000 rabbit-α-ERD2 (BEI MRA-1). Blots were washed 3 times in TBST and were incubated for 1 hour at room temperature in a 1:10,000 dilution of an appropriate fluorescent secondary antibody (Li-COR Biosciences). Blots were washed 3 times in TBST and once in PBS before imaging on a Li-COR Odyssey imager.

### Actinonin treatment and IPP rescue

To generate apicoplast-minus parasites, ring-stage cultures were treated with 10 μM Actinonin (Sigma A6671) and 200 μM isopentenyl pyrophosphate (IPP, Isoprenoids LLC) for 3 days.

### Protease protection assay

50 mL of schizont stage parasite culture at 2% hematocrit and ~5% parasitemia was separated from erythrocytes by lysis in 0.1% saponin for 5 minutes on ice. Parasite pellets were washed 3 times with ice-cold PBS and lysed in hypotonic lysis buffer (20 mM HEPES-NaOH, pH 7.2) by passing through a 27-gauge needle 30 times. Lysate was centrifuged in assay buffer (200 mM HEPES-NaOH pH 7.4, 200 mM NaCl, 1 M sucrose) 3 times for 10 minutes at 1,500 x g to remove unbroken cells and debris. Lysate was split into 4 samples: (i) no treatment (ii) 50 μg/mL Proteinase K (Thermo Fisher EO0491) (iii) 1% Triton X-100 (Sigma 45ZH27) (iv) 50 μg/mL Proteinase K and 1% Triton X-100. After incubating for 30 minutes on ice, proteolysis was stopped by addition of 1 mM phenylmethylsulfonyl fluoride (PMSF, Sigma P7626). Samples were analyzed by western blotting as described. Band intensities were quantified using Image Studio Lite software (Li-COR Biosciences).

### Knockdown growth assays

Ring-stage TetR-DOZI strain parasites were washed twice in culture medium to remove ATc. Parasites were divided into 3 cultures supplemented with 500 nM ATc, no ATc, or no ATc + 200 μM IPP. Samples were collected at the schizont stage of each growth cycle for flow cytometry analysis and western blot. Parasites in each condition were diluted equally into fresh medium with 2% hematocrit for 4 reinvasion cycles. For parasitemia measurements, aliquots of culture were incubated with 16.67 μM dihydroethidium (Thermo Fisher) for 15 minutes. Parasites were analyzed on a BD Accuri C6 flow cytometer and 100,000 events were recorded for each condition. Two-way ANOVAs were performed in GraphPad Prism. For western blots, equal volumes of culture were loaded for each treatment condition of a strain.

### Immunofluorescence microscopy

Parasite cultures were washed once in PBS and were fixed with 4% paraformaldehyde (Electron Microscopy Sciences 15710) and 0.0075% glutaraldehyde (Electron Microscopy Sciences 16019) in PBS for 20 minutes. Cells were washed once in PBS and allowed to settle onto poly-L-lysine-coated coverslips (Corning) for 60 minutes. Coverslips were washed once with PBS, permeabilized in 0.1% Triton X-100 in PBS for 10 minutes, and washed twice more in PBS. Cells were treated with 0.1 mg/mL NaBH_4_/PBS for 10 minutes, washed once in PBS, and blocked in 5% bovine serum albumin (BSA)/PBS. Coverslips were incubated overnight at 4°C in primary antibody diluted in 5% BSA/PBS using the following concentrations: 1:500 rabbit-α- *Pf*ACP^48^ (kind gift from S. Prigge) and 1:100 rat-α-HA 3F10 (Sigma 11867423001). Coverslips were washed 3 times in PBS, incubated with secondary antibodies donkey-α-rabbit 568 (Thermo Fisher A10042) and goat-α-rat 488 (Thermo Fisher A-11006) at 1:3,000 dilution, and washed 3 times in PBS. Coverslips were mounted onto slides with ProLong Gold antifade reagent with DAPI (Thermo Fisher) and were sealed with nail polish prior to imaging.

Cells were imaged with 100X, 1.4 NA or 100X, 1.35 NA objectives on an Olympus IX70 microscope with a DeltaVision system (Applied Precision) controlled with SoftWorx version 4.1.0 and equipped with a CoolSnap-HQ CCD camera (Photometrics). Images were captured as a series of z-stacks separated by 0.2 μm intervals and displayed as maximum intensity projections. Brightness and contrast were adjusted using Fiji (ImageJ) for display purposes.

### Yeast transformation and halo assay

Transformation was performed as described previously^49^. MFA1 expression plasmid along with complementation constructs expressing *Sc*Rce1, AMR4^112-432^, AMR4^197-432^, or empty vector were transformed into either AFC1 RCE1 (JRY5460), *afc1Δ* RCE1 (JRY6095), AFC1 *rce1Δ* (JRY5462), or *afc1Δ rce1Δ* (JRY5463) *S. cerevisiae* strains^32^.

For halo assays^32,50^, cultures of Matα *sst2* (JRY3343) and transformed MAT**a** cells were grown overnight. 4×10^6^ MATα *sst2* cells were spread onto prewarmed YPD + 0.04% Triton X-100 plates and allowed to dry. 1×10^6^ MAT**a** cells in 5 uL were then spotted onto the lawn and allowed to dry. Plates were incubated at 30°C 2-3 days before imaging. Image color was inverted and brightness and contrast were adjusted in Fiji (ImageJ) for display purposes.

### Sequence analysis and phylogeny

Protein sequences were retrieved from EuPathDB^51^ and UniProt^52^. The CPBP domain of each gene was annotated using InterProScan^53^. Sequences were aligned using MAFFT^54^ and unrooted maximum likelihood phylogenies were performed using IQ-TREE^55^ with default settings and 100 standard bootstrap intervals. Phylogenetic trees were then imported into FigTree (v.1.4.4) and midpoint rooted. Targeting sequences for each protein were predicted using TargetP-2.0^56^ (Table S2).

### Whole-genome sequencing and SNP analysis

Isolation of apicoplast-minus mutant clones and DNA extraction were described previously^14^. Genomic DNA samples from 6 additional clones were used to generate paired-end libraries using the NEB Next Ultra FS II kit with 8 PCR cycles. Libraries were sequenced on an Illumina NextSeq 500 using 2×150 bp paired-end sequencing (Chan Zuckerburg Biohub). Single nucleotide polymorphism (SNP) analysis was performed using a custom script as described previously^14^. Briefly, adapters were removed using cutadapt, and sequencing reads were aligned to the *P. falciparum* 3D7 (version 35) genome using Bowtie2. PCR duplicates were removed and raw SNP’s (base quality ≥ 20) were called using Samtools. Bcftools was used to generate the raw variant list for parental (allele frequency ≥ 0.01, depth ≥ 1) and mutant (allele frequency ≥ 0.9, depth ≥ 20) strains. Variants were filtered to only include protein-coding mutations, and variants found in the parental strain were subtracted from the mutant variant list. Mutations in hypervariable gene families and mutations that were previously annotated in PlasmoDB were excluded from the final variant list (Table S1).

## Acknowledgements

We thank Sean Prigge for the α-ACP antibody, Walid Houry for the α-ClpP and α-ClpR antibodies, Sebastian Mikolajczak and Stefan Kappe for the α-*Py*BiP antibody, and Jacquin Niles for the NF54^Cas9+T7 Polymerase^ strain and pSN054-V5 plasmid.

This work was supported by the following funding sources: NIH 1R01AI141366 (EY), Burroughs Wellcome Fund (EY), Chan Zuckerberg Biohub (EY), NIH T32GM007276 (TRM), and NIH T32GM007365 (YT). The funders had no role in study design, data collection and interpretation, or the decision to submit the work for publication.

