## Supplemental Material for "CaaX-like protease of cyanobacterial origin is required for complex plastid biogenesis in malaria parasites"

**Supplemental Materials:**

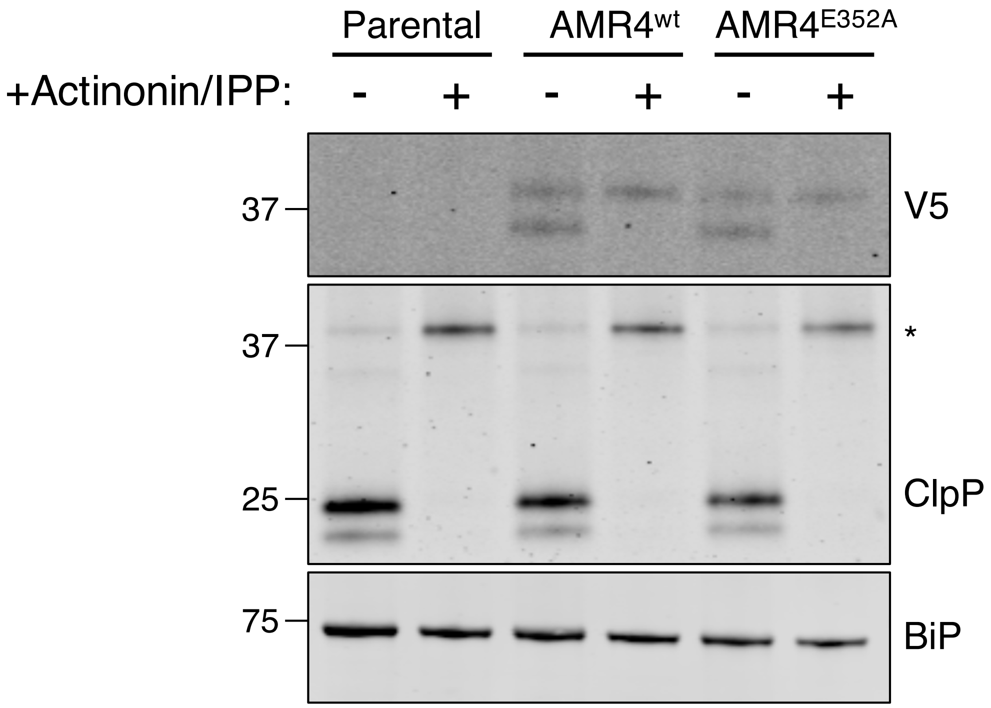


**Figure S1. Episomally expressed AMR4$^{wt}$ and AMR4$^{E352A}$ are imported into the apicoplast.** Western blot showing expression and processing of V5-tagged AMR4$^{wt}$ and AMR4$^{E352A}$ after 3 days of either no treatment or +10 µM actinonin/200 µM IPP to eliminate the apicoplast. AMR4$^{wt}$ and AMR4$^{E352A}$ are both properly imported into the apicoplast, as demonstrated by their apicoplast-dependent transit peptide cleavage. ClpP serves as a positive control for apicoplast import, and BiP serves as a loading control. Parental strain is AMR4 TetR-DOZI without episomal complementation.

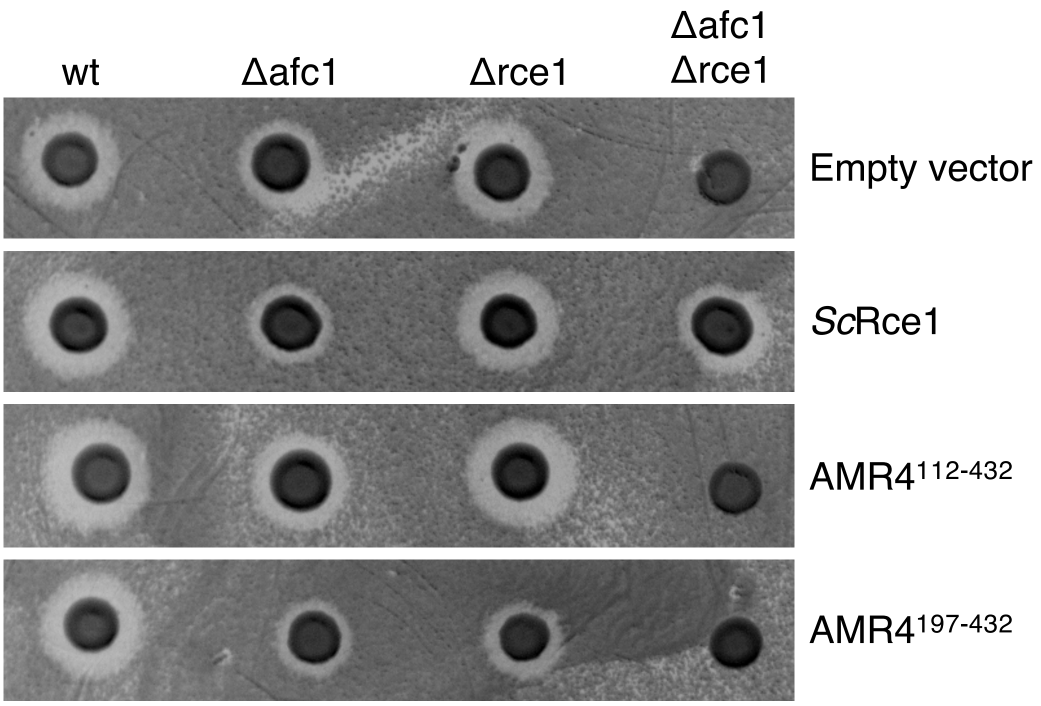


**Figure S2. AMR4 does not functionally complement CaaX proteases in yeast "halo" assay.**

Proper secretion of the yeast mating pheromone **a**-factor is dependent on CaaX prenylation and post-prenylation processing. A mutant α-type strain (MATα$^{sst2}$) is growth arrested when exposed to secreted **a**-factor, such that when MAT**a** cells are spotted over a lawn of MATα$^{sst2}$ cells, proper CaaX processing of **a**-factor results in a growth arrest "halo" around the spot [Chan 1982, https://doi.org/10.1128/MCB.2.1.11]. This assay has been used previously to test heterologous CaaX proteases, and is sensitive enough to detect below 5% of wild-type **a**-factor processing [Cadiñanos 2003a, https://doi.org/10.1074/jbc.M306700200; Cadiñanos 2003b, https://doi.org/10.1042/bj20021514; Michaelis 2012, https://doi.org/10.1128/MMBR.00010-12]. We genetically modified MAT**a** cells and used this halo assay as a read-out for functional CaaX protease activity. Double knockout of both yeast CaaX proteases, Rce1 and Afc1, abolished the halo phenotype as previously reported [Trueblood 2000, https://doi.org/10.1128/MCB.20.12.4381-4392.2000]. Complementing this strain with *Sc*Rce1, which contains a CPBP domain, largely rescues the halo phenotype indicating a restoration of CaaX post-prenylation processing. We tested two AMR4 constructs, containing either the post-transit peptide protein (AMR4$^{112-432}$) or the Rce1-like domain only (AMR4$^{197-432}$), each fused with the N-terminal *Sc*Rce1 signal peptide for proper targeting. Both constructs failed to rescue the halo phenotype, however we were unable to detect the tagged AMR4 constructs by western blot and therefore cannot confirm proper protein expression or localization. Images are representative of two independent experiments with two technical replicates per experiment.

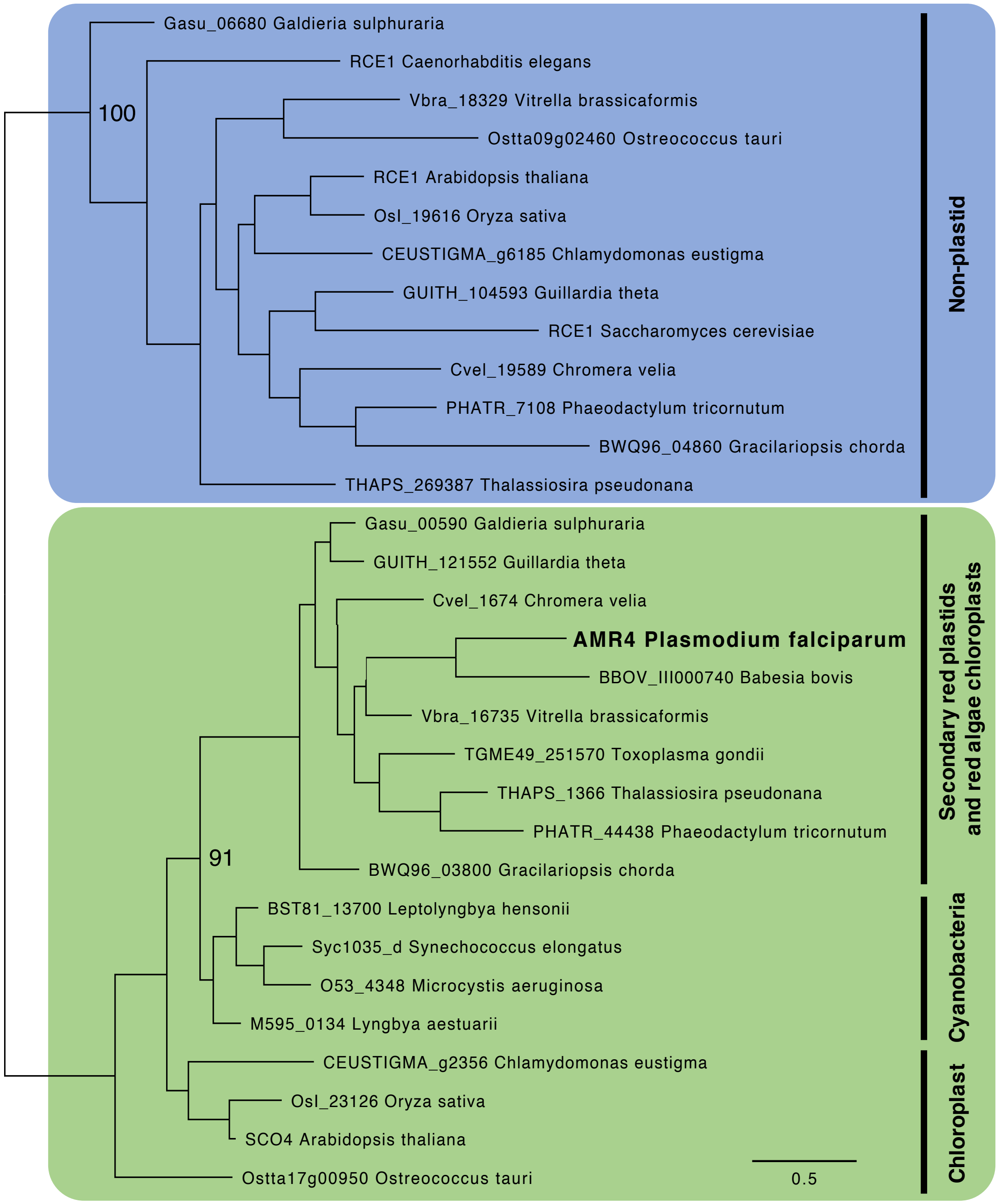


**Figure S3: Phylogenetic analysis of CPBP domains confirms cyanobacterial origin of AMR4.**

Phylogenetic analysis of selected CPBP proteins from cyanobacteria, primary and secondary plastids, and non-plastid Rce1's. To prevent bias from different targeting sequences, only the annotated CPBP domain (IPR003675) from each protein was used. Maximum likelihood phylogeny defines a monophyletic clade of cyanobacterial and chloroplast-targeted proteins along with AMR4 and its secondary plastid homologs. Outside of this clade are non-plastid targeted Rce1 proteins from eukaryotes with and without plastids. Branch support values for well-supported major nodes are shown, out of 100 bootstrap intervals.

**Table S1:** Raw nucleotide variants identified in sequenced clones.

**Table S2:** List of proteins and predicted localizations used for phylogenetic analysis.

**Table S3:** Primer and gBlock sequences used in this study.
